# Validation of a Lysis Buffer Containing 4 M Guanidinium Thiocyanate (GITC)/ Triton X-100 for Extraction of SARS-CoV-2 RNA for COVID-19 Testing: Comparison of Formulated Lysis Buffers Containing 4 to 6 M GITC, Roche External Lysis Buffer and Qiagen RTL Lysis Buffer

**DOI:** 10.1101/2020.04.05.026435

**Authors:** Martina F. Scallan, Catherine Dempsey, John MacSharry, Isabelle O’Callaghan, Paula M. O’Connor, Conor P. Horgan, Edel Durack, Paul D. Cotter, Sarah Hudson, Humphrey A. Moynihan, Brigid Lucey

## Abstract

The COVID-19 pandemic has resulted in increased need for diagnostic testing using reverse transcriptase real-time PCR (RT-PCR). An exponential increase in demand has resulted in a shortage of numerous reagents in particular those associated with the lysis buffer required to extract the viral RNA. Herein, we describe a rapid collective effort by hospital laboratory scientists, academic researchers and the biopharma industry to generate a validated lysis buffer. We have formulated a 4M Guanidinium thiocyanate (GITC)/ Triton X-100 Lysis buffer which provides comparable results with the recommended reagents. This buffer will ease the burden on hospital labs in their heroic efforts to diagnose a large population of patients.

## Introduction

The global demand for reagents for real-time reverse transcriptase PCR (RT-PCR) diagnostic tests for COVID-19 has caused a bottle neck across Ireland and the UK in efforts to follow the World Health Organisation’s advice to “*Test, Test, Test”*.

Expansion of PCR testing is critical to gain control over the pandemic spread of COVID-19 and the world-wide shortage of lysis buffer is a rate limiting step. Here we disseminate details of a lysis buffer formulation that came into existence as a consequence of a rapidly formed multi-centre collaboration. A replacement lysis buffer for extraction of viral nucleic acids from respiratory samples was desperately needed by diagnostic laboratories. A panel of new lysis buffer formulations was generated and validated for use in COVID-19 testing by virus-specific RT-PCR and are being distributed throughout Ireland. It is hoped that by sharing the formulation and validation, we can make it useful to others in the national and international scientific community.

The key chemical constituent of the lysis buffer (guanidinium thiocyanate) has become scarce, scientists from the School of Microbiology, University College Cork, combined their long-held knowledge of lysis buffers for RNA extraction, with insights from published work relating to extraction of RNA using magnetic glass beads (Chomczynski & Sacchi, 1987; Hui He *et al*., 2017). Ultimately, they formulated a lysis buffer, which uses much less guanidinium thiocyanate than standard formulations appear to use (4 M compared to 6 M). Simultaneously, scientists at Teagasc, Fermoy were working on a different experimental formulation for lysis buffer (Boom *et al*., 1990). Four laboratory preparations in total were tested for efficacy by medical scientists based in the Clinical Microbiology Department at Cork University Hospital (CUH), who conducted validation experiments using positive and negative samples. After validation, pharma-based scientists, generously prepared a large volume of the much needed 4 M GITC lysis buffer for distribution by CUH.

### Virology and RNA biology expertise

Dr Martina Scallan & Dr John Mac Sharry

### Buffer preparation

Dr John Mac Sharry & Dr Martina Scallan; Prof Paul Cotter & Paula O’Connor

### Evaluation of buffer preparations and validation for use for in-vitro diagnosis of patients tested for Covid-19

Senior medical scientists Catherine Dempsey & Isabelle O’Callaghan

### Risk Assessment for buffer preparation

Dr Edel Durack & Dr Sarah Hudson

### Co-ordination of preparation of 4.5 L of the 4 M formulation for distribution by CUH

Dr Humphrey Moynihan & Dr Conor Horgan

### Co-ordination/preparation of paper

Dr Brigid Lucey

**It should be noted that the following chemical preparation, after validation was selected for use with named commercial platforms and testing kits. We recommend that each centre adopting it in their laboratories should first validate it for use with their own systems. It is also advisable to provide a lot number with each batch prepared for traceability**.

The 4 M GITC lysis buffer was developed based on knowledge of the role of GITC in RNA extraction and protection (Chomczynski & Sacchi, 1987; Boom et al., 1990), adapted for magnetic glass bead extraction (Hui He *et al*., 2017), and validated against recognised standards using positive and negative controls. Details of the composition of 4 M GITC lysis buffer and for the preparation of 1 litre are given below.

The lysis procedure used in the diagnostic laboratory mixes lysis buffer and sample 1:1, this generates a working concentration of 2 M GITC during the lysis step as reported to be optimal by Hui He *et al*., 2017.

**Buffer Composition:**

4 M guanidinium thiocyanate (GITC)

55 mM* Tris-HCl

25 mM EDTA (Ethylenediaminetetraacetic acid)

3 % (v/v) Triton X-100

0.01 % (w/v) Bromophenol blue

(*NOTE: calculated from the total amount of 0.1 M Tris pH 7.6 added, diluted by the degree of volume expansion observed when the GITC goes into solution).

**Preparation of one litre of 4M Guanidinium thiocyanate (GITC)/ Triton X-100 Lysis buffer** <u>(please also read risk assessment):</u>

1. 472.75 g of GITC is brought into solution initially by adding 400 ml of 0.1 M Tris HCl Ph 7.6. This will require heating in a 65°C water bath and some shaking of the vessel (but with lid well secured). In our hands, once fully dissolved the volume of the solution was 600 ml.
2. Make up to 750 ml with 0.1 M Tris HCl pH 7.6.
3. Add 50 ml of 0.5 M EDTA, mix.
4. Add 30 ml Triton-X-100, mix
5. Volume made up to 1 L with 0.04 % (w/v) bromophenol blue (DEPC-treated water can be used instead)

**Notes:**

- See Appendix 1 for chemical agent risk assessment sheet
- Use laboratory coat, safety goggles, gloves, chemical respirator dust mask (N95 mask)
- Open containers of guanidinium thiocyanate powder should be handled in a fume cabinet
- The balance for weighing should be positioned outside the fume cabinet (but close by). Receiving vessel must have a lid that can be sealed for weighing
- Significant chemical or physical hazards notable when preparing buffer: poisoning and environmental hazards
- Buffers made up with DEPC-treated molecular biology grade water
- A significant increase in volume is observed upon GITC dissolution and the degree of volume expansion was found to vary between preparations performed at different sites. This may reflect variations related to the manufacture of GITC and/or storage conditions of the chemical and accentuates the need for individual site validation before use diagnostically.

Validation of the preparation was conducted using two different automated extraction instruments and two different detection instruments.

The validation consisted of a comparison of the efficacy of extraction with four laboratory-prepared buffers and both a Qiagen (RLT lysis buffer from an RNeasy kit Cat. No./ID: 79216) and a Roche external lysis buffer when testing positive (amplifying two Covid-19 specific targets) and negative controls. (In each case the extraction volumes were 200 μl of buffer:200 μl transport medium.) Both positive and negative controls incorporated an internal control. Detection of targets was compared for crossing point values during real time Polymerase Chain Reactions (RT PCR).

## Results

Detection of targets was compared for crossing point values (Cp) for the four prepared buffers and the Roche external lysis buffer as shown in **Table 1**. Automated extraction was conducted using the Roche MagNa pure LC and the PCR was performed on the Roche z480 RT PCR instrument. On a second day, the 4 M GITC buffer was tested in comparison with Roche and Qiagen buffers, using the IndiMag48 extraction system and RT PCR was conducted on a Roche LightCycler II 480 (**Table 2**). On the basis of these combined data, the buffer containing 4 M GITC was confirmed as suitable for use with the diagnostic systems in CUH. **Figure 1** shows the relative performance of all buffers tested, measured using Cp values Based on these data, the 4 M GITC lysis buffer was selected for future preparation and use when extracting respiratory samples for Covid-19.

**Table 1.** Comparison of the efficacy of four formulations of lysis buffer intended for the extraction of viral RNA from respiratory samples for Covid-19 testing when compared with the Roche external lysis buffer

| Sample type | SARS CoV2 Cp<sup>^</sup> | B beta CoV Cp<sup>^</sup> | Internal Control Cp<sup>^</sup> | Lysis Buffer |
| --- | --- | --- | --- | --- |
| Positive sample | 22.37 | 22.96 | 29.46 | 4 M GITC (pH7.6) with Triton |
| Negative sample | Not Detected | Not Detected | 31.42 | 4 M GITC (pH7.6) with Triton |
| Positive sample | 22.36 | 23.36 | 29.62 | 5.4 M GITC (pH6.4) with Triton |
| Negative sample | Not Detected | Not Detected | 31.18 | 5.4 M GITC (pH6.4) with Triton |
| Positive sample | 22.29 | 23.25 | 29.44 | 6 M GITC (pH7.6) with Triton |
| Negative sample | Not Detected | Not Detected | 32.98 | 6 M GITC (pH7.6) with Triton |
| Positive sample | 22.57 | 21.69 | 28.82 | 4.75 M GITC (pH 7.6) without Triton |
| Negative sample | Not Detected | Not Detected | 30.76 | 4.75 M GITC (pH 7.6) without Triton |
| Positive sample | 20.85 | 21.62 | 27.32 | Roche Lysis buffer |
| Negative sample | Not Detected | Not Detected | 28.59 | Roche Lysis buffer |

<sup>^</sup>Cp = Crossing point during the test reaction cycles at which test is denoted positive; The complete formulations for the 5.4 M, 6 M GITC preparations with Triton X-100 and the 4.75 M GITC formulation without Triton are not currently listed. pH values refer to the pH of 0.1 M Tris-HCl used in buffer preparation.

**Table 2.**
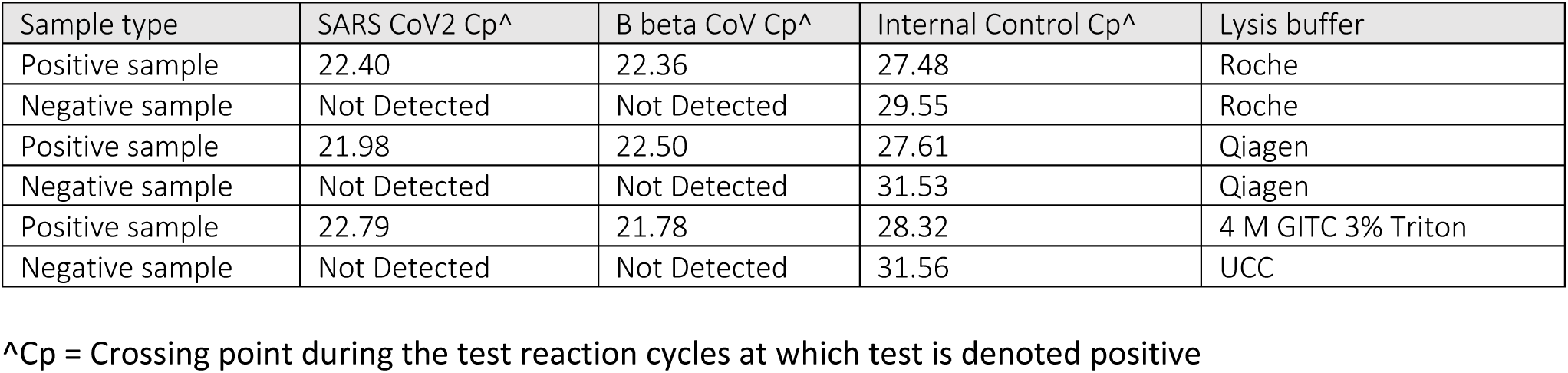
Comparison of the efficacy of 4 M GITC (with 3% Triton X-100) with Qiagen and Roche lysis buffers

**Figure 1.**
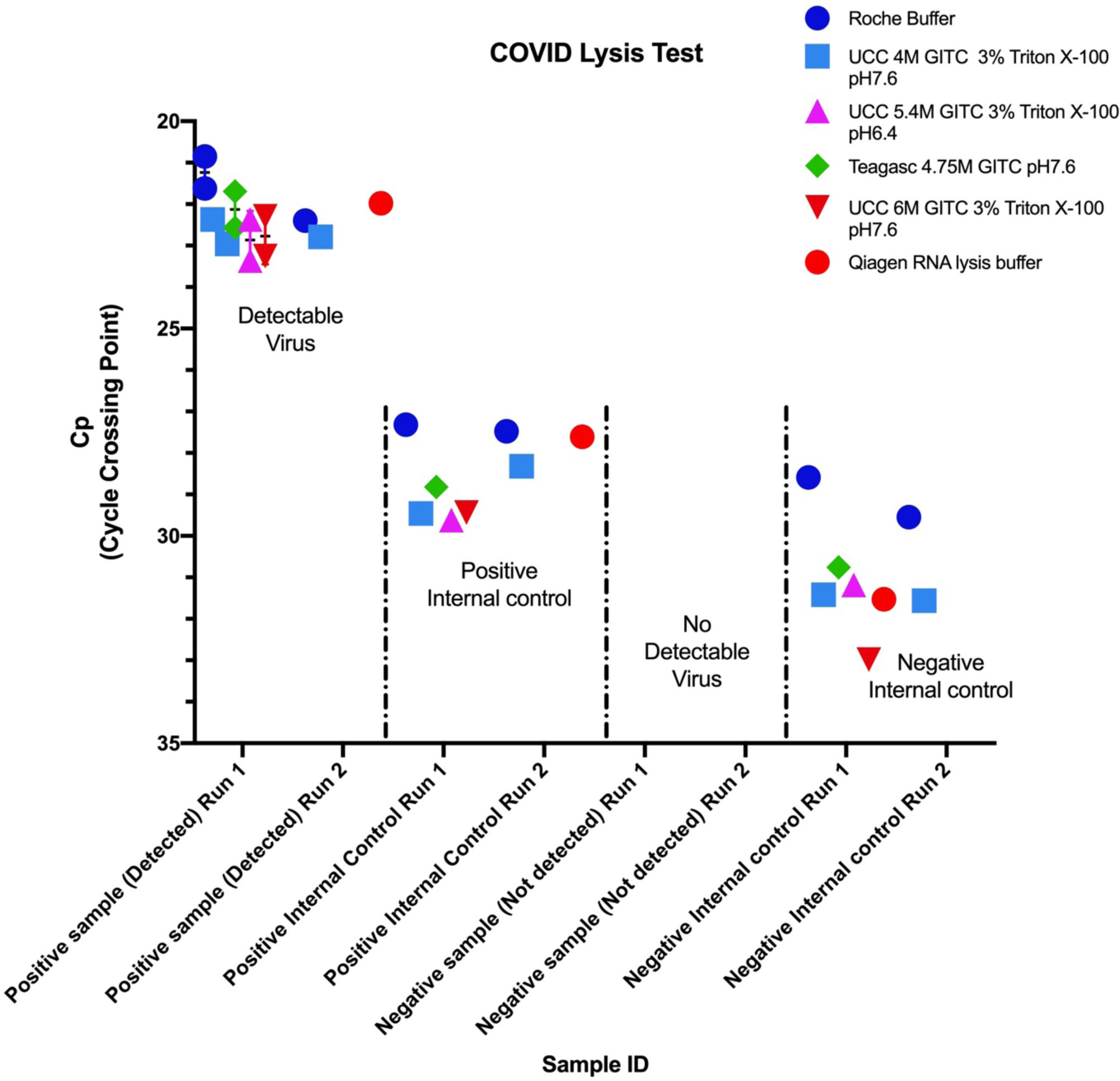
Comparison of the performance of commercial and non-commercial buffers as indicated by Cp values (y-axis) when tested against positive and negative sample controls during real-time RT-PCR diagnostic test for Covid-19.

## Discussion

A 4 M GITC lysis buffer with 3 % (v/v) Triton X-100 has been shown to work very well in COVID-19 RT-PCR, it includes a detergent which helps to disintegrate the virus during extraction, bromophenol blue as a visual aid for addition of lysis buffer to clinical samples and has the advantage of using less of a scarce ingredient. The knowledge that 4 M guanidinium thiocyanate (GITC) is sufficient in this lysis buffer means the straitened global supply of this chemical can be shared more effectively to make more lysis buffer to perform more tests around the world and help the world to take back control against this pandemic virus.

Heeding Dr Michael J. Ryan’s advice “Perfection is the enemy of the good when it comes to emergency management”, we do not want to delay in communicating our findings to the wider world. Further variations of the 4 M GITC lysis buffer formulation can be tested further, in particular formulations with graded increases in % w/v of Triton X-100 could be compared. Work is ongoing to test if variations in the molarity of Tris-HCl affect lysis buffer performance. Of highest priority is to get the message out that this 4 M GITC lysis buffer works well. In addition we observed that Qiagen’s RLT lysis buffer from RNeasy kits worked very well with the systems and processes in place in the diagnostic laboratory at Cork University Hospital.

When Covid-19 testing began, the individual diagnostic microbiology hospital laboratories in Ireland agreed not to stockpile any of their resources from any of the other Irish hospital laboratories. Through their professional body, the Academy of Clinical Science and Laboratory Medicine, they had daily meetings (using WhatsApp) to problem solve as a group where necessary.

Normally, hospital laboratories buy in reagents that allow them to concentrate on the business of diagnosis of patients and the preparation of their own reagents is rare. However, a matter of particular concern was the unexpected short supply of one reagent (lysis buffer) needed to lyse the virus causing COVID-19, to facilitate RNA extractions and subsequent detection. In straitened times, society needs to pool its resources and expertise to best impact and this case study and validation represents a generous and rapid response from the scientific community, when called upon by the Academy of Clinical and Laboratory Medicine, at a time when large international commercial companies were unable to supply this vital reagent.

In the midst of this combined effort, research and medical scientists in other parts of Ireland are communicating their validation results in turn. Part of the group effort in the case being described here is that of the two scientists in the University of Limerick, who prepared the risk assessment documentation for the report, while simultaneously making up buffer and supplying their nearest hospital laboratory at University Hospital Limerick.

In an urgent situation, Irish scientists, academics and biopharma have worked collectively to pool expertise, technical know-how, chemicals from their research laboratories, their facilities, time, co-ordination and communication skills along with determination, to develop a suitable replacement lysis buffer in less than two weeks. Like hospital laboratories, third level research laboratories rely on buying ready-to-go kits and reagents to speed up their research. Fortunately, retention of the knowedge regarding the chemical composition of the buffers among the scientists allowed them to make up their own reagents from raw materials.

## Acknowledgements

The authors wish to acknowledge colleagues in Sligo Institute of Technology and at University College Hospital, Limerick in the Departments of Serology and Microbiology for their valuable assistance during consideration of lysis buffer formulations. Also, Dr John O’Callaghan, Senior Technical Officer, School of Microbiology, UCC for technical assistance during the UCC stages of development and preparation (and for providing the Qiagen buffer), Dr Ewen Mullins of Teagasc for sourcing and sharing reagents, and Dr Darren Crowley and Dr Eoghan Casey who were involved in the 4.5 L buffer preparation at Eli Lilly.

## Appendix 1. Risk Assessment of agents used in the preparation of lysis buffer containing Guanidine Thiocyanate, Trizma Base, EDTA disodium salt, Triton-X and Bromophenol Blue

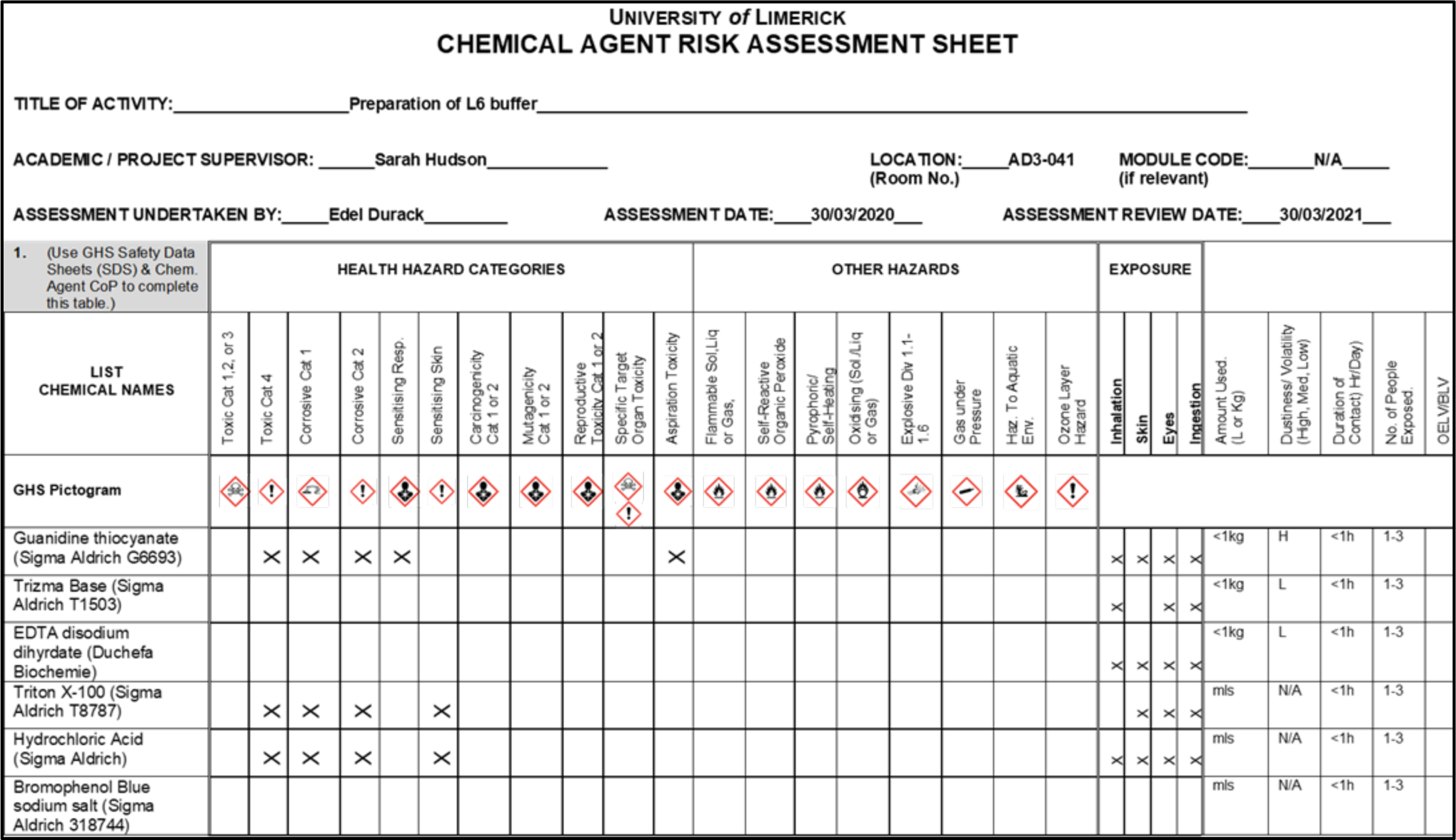

